# Fluid flow induced biomechanical origin of collagen architecture in articular cartilage

**DOI:** 10.1101/2025.07.13.664559

**Authors:** Dhruba Jyoti Mech, Mohd Suhail Rizvi

## Abstract

The zonal collagen architecture of articular cartilage (AC) is essential for its mechanical function and long-term homeostasis. While its structural organization is well established, the mechanistic basis for the emergence and maintenance of this architecture remains unresolved. In this study, we propose a fluid flow–driven mechanism for the evolution of collagen fiber orientation in AC, using both a continuum orientation field model and a discrete three-dimensional fiber network model. Joint movements, shear-dominated during embryogenesis and combined shear-compression postnatally, induce synovial fluid flow, which guides collagen alignment through preferential fiber deposition. Our models reproduce the characteristic Benninghoff architecture observed in mature AC and are validated against experimental data across multiple species, joint types, and developmental stages. We demonstrate how joint- and organism-specific mechanical loading leads to diverse collagen arrangements and zonal organization. Further, by systematically varying shear and compressive loading durations to mimic different physical activities, we show that the collagen architecture, mechanical stiffness, and effective synovial fluid viscosity of AC adapt in an activity-dependent manner. Finally, we simulate osteoarthritic remodeling as a localized disruption to fluid flow and show how it leads to progressive collagen disorganization. These findings offer a unifying biomechanical framework for AC development, function, and degeneration, with implications for tissue engineering and rehabilitation strategies.

## 1 Introduction

Articular cartilage (AC), the specialized connective tissue covering the ends of bones in synovial joints, provides a smooth, low-friction surface for movement and distributing mechanical loads to protect underlying bone (Smith et al., 2019). It is composed primarily of water, collagen (predominantly type II), proteoglycans, and chondrocytes, which together form a highly organized extracellular matrix optimized for mechanical function and durability (Bhosale and Richardson, 2008; Carballo et al., 2017; Semitela et al., 2024). The collagen architecture in articular cartilage is a multilayered structure, usually stratified into three zones- the superficial zone, where collagen fibers are tightly packed and aligned parallel to the articular surface; the middle zone, characterized by a relatively random fiber orientation; and the deep zone, where fibers are oriented perpendicularly and anchor into the subchondral bone (Bhosale and Richardson, 2008; Carballo et al., 2017). This mature zonal organization emerges progressively during embryonic and post-natal development.

In neonatal articular cartilage, the collagen arrangement is relatively simple with collagen fibers predominantly oriented parallel to the articular surface (in the superficial layer) or arranged in random orientations (in deep layers) (Clark et al., 1997). As the cartilage matures during post-natal stages, the collagen network undergoes significant remodeling to develop the zonal structure and biomechanical properties characteristic of adult articular cartilage (Decker et al., 2015; Decker, 2017). In sheep, the articular cartilage collagen remodels from a parallel network at birth to a Benninghoff network by maturity (van Turnhout et al., 2010). In particular, during postnatal development, collagen remodeling begins just below the future transitional zone. Similarly, equine studies also show collagen in articular cartilage remodels from parallel to Benninghoff architecture within months after birth, with collagen’s predominant orientation being parallel to the articular surface throughout the entire cartilage depth (van Turnhout et al., 2008; Cluzel et al., 2013). A recent <u>in vitro</u> study using chondrocyte-derived cartilage organoids has shown the evolution of collagen arrangement (Peters et al., 2024). It has also been suggested that during postnatal development AC’s structural maturation is driven by tissue resorption and neoformation rather than internal remodeling (Hunziker et al., 2007).

Despite the prevalence of the three-layered collagen fiber architecture of AC in literature, it is not uniform across all joints but varies significantly depending on joint-type. As shown in (He et al., 2013) the collagen arrangements in the distal humerus and femoral condyle are vastly different in kangaroo. In the femoral condyle, the articular cartilage has the characteristic three-layered Benninghoff architecture whereas in the distal humerus, the cartilage appeared fine with no clear zonal organization. These joint-type and developmental stage dependent differences in collagen architecture suggest that local mechanical loading environments may influence the structural organization of articular cartilage.

During degenerative diseases, such as osteoarthritis (OA), articular cartilage, due to its lack of regeneration capabilities, undergoes progressive degradation, including collagen network disruption, leading to impaired mechanical properties and joint degeneration (Wieland et al., 2005; Bhosale and Richardson, 2008). With disease progression, the collagen arrangement in intact articular cartilage also undergoes structural changes, such as human patellar cartilage showing changes in collagen fiber orientation and dispersion with advanced OA (Saarakkala et al., 2010). Currently, no therapy effectively halts the structural degradation of cartilage and bone or reverses existing structural defects caused by osteoarthritis. In the past couple of decades, there have been numerous works on AC regeneration with the help of tissue engineering approaches by combining biomaterials, cells, and biochemical and biophysical factors to create functional cartilage tissue (Becerra et al., 2010; Semitela et al., 2024). These works have identified and optimized the scaffold material and architecture, cells, and biochemical factors. However, the biomechanical stimuli required for AC regeneration have been elusive. Insights into the role of biomechanics in embryonic and post-natal articular cartilage development can provide critical guidance for AC tissue engineering.

There have been several works acknowledging the role of mechanical loading of the joints on the overall development of the collagen arrangement (Fell and Canti, 1934; Persson, 1983; Hyttinen et al., 2009; Brama et al., 2009, 2000; Rieppo et al., 2009; Julkunen et al., 2010). For embryonic development, it has been shown that limb paralysis during embryogenesis disrupts or prevents joint formation, presumably for maintaining cavitation and synovial fluid formation (Osborne et al., 2002). However, the exact mechanistic link between mechanical stimulation and the structural organization of articular cartilage remains elusive.

In this work, we propose a biomechanical mechanism underlying the development of collagen architecture in articular cartilage. Specifically, we consider the nature of the synovial fluid flows during the late stages of embryonic and post-natal development and growth and propose its central role in the evolution of collagen architecture in articular cartilage during normal development, physiology, and disease conditions, such as osteoarthritis.

## 2 Mathematical model

### 2.1 Basic assumptions

We consider articular cartilage to be a porous medium with permeability that depends on the underlying collagen fiber arrangement (Fig. 1A). In the absence of collagen fibers or with isotropic collagen fiber distribution, we consider articular cartilage permeability to be isotropic. This assumption is justified by the isotropic nature of the base matrix. The interface between AC and subchondral bone is assumed to be permeable to the exchange of fluids and solutes. This assumption is supported by the presence of microchannels and vascular networks in subchondral bone (Clark, 1990; Taheri et al., 2019). Even though the synovial fluid has been known to be non-Newtonian, here we consider it to be a Newtonian fluid with a high viscosity (approximately of the order of 10 mPa-s, (Fu et al., 2019)) that plays a key role in joint lubrication and load-bearing.

**Figure 1:**
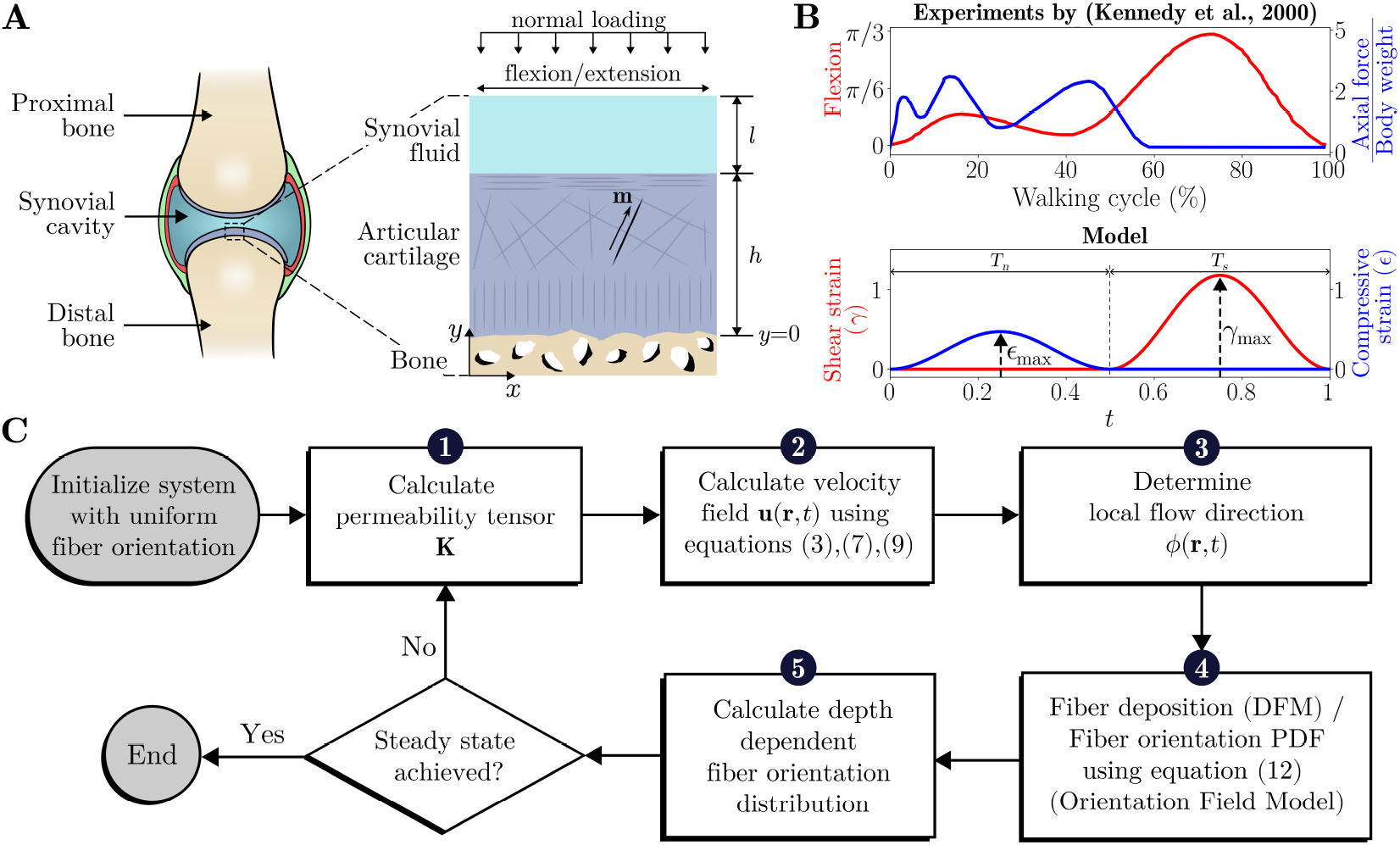
An overview of the modeling framework and analysis. (A) Schematic representation of a typical joint anatomy. The magnified section highlights the region selected for modeling in this work. (B) Comparison of loading cycles used in the model with experimentally observed loading patterns (data taken from (Kennedy et al., 2000)) during a typical walk cycle.(C) Flowchart illustrating the numerical simulation workflow for the coupled model of articular cartilage mechanics and collagen fiber remodeling. Both the orientational field model and the discrete fiber model follow this common simulation framework.

In this work, we are focusing on the changes in the collagen fiber architecture, and we assume that the articular cartilage volume does not change over time. It is to be noted that this is a simplification and is taken to remove the effect of the volumetric growth of AC on collagen arrangement. In a more general modeling of the process, one has to incorporate AC growth, which can be significant in the late stages of embryonic development and early post-natal stages (Hunziker et al., 2007). During these stages of AC development and remodeling, we assume collagen synthesis by chondrocytes and that its degradation follows first-order kinetics. We also make the simplifying assumption that chondrocyte density is uniform across the depth of the cartilage.

During embryonic development, joint movements predominantly generate shear stresses on the developing articular cartilage (AC), as evidenced by experimental observations of coordinated flexor and extensor muscle activity (Chambers et al., 1995; Bradley et al., 2005; Ryu and Bradley, 2009). However, post-natal development introduces more complex loading patterns, but for simplicity, these can be approximated as a superposition of normal and shear stresses, reflecting the combined effects of weight-bearing and joint movement which similar to the experimental measurements (top panel in Fig. 1B, see (Kennedy et al., 2000) for details). Therefore, we assume that the loading pattern is shear loads during embryonic development and a combination of normal and shear loads <u>post-natum</u> (bottom panel in Fig. 1B).

### 2.2 Model formulation

We model the mechanics of a small portion of articular cartilage that spans its thickness (schematic in Fig. 1A), and we ignore the curvature in the articulating surfaces. As shown in Fig. 1A, the *x*-*y* plane is normal to the articulating surface with *x*-direction aligning with the direction of relative AC motion during joint movement. Furthermore, to keep the model simple, we also assume that the dependence of any property along the third direction *z* is minimal. The domain of interest in the model is taken to be one half of the articulating joint (as shown in the enlarged version in Fig. 1A) and it is considered to be of layered arrangment of bone, articular cartilage (thickness *h*) and synovial cavity (thickness *l*). The proposed model can easily be extended to its more general form by considering the curvature in AC and the details of the three-dimensionality of the joint movements.

#### Joint movement driven synovial fluid flow

The structure of synovial joints motivates us to model the fluid flow between the two articulating ends of the joints as the flow between two parallel plates. Due to the porous nature of AC, it requires the model to incorporate this information. As shown in the schematic (enlarged section of Fig. 1A), we consider one-half of the section of the articulating joint, which is formed by two zones- the synovial cavity and articular cartilage. In the synovial cavity, the fluid flow can be described by the Navier-Stokes equations

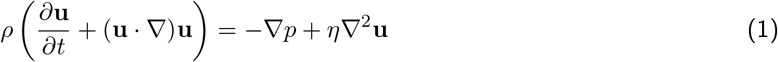

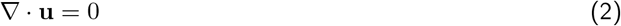

where *ρ* is the fluid density, **u** is the velocity field, *p* is the pressure, and *η* is the dynamic viscosity of the synovial fluid. In order to keep the model simple, we will drop the nonlinear term (**u** · ∇)**u**, but this simplification can be relaxed without changing the overall conclusions of this work. In the porous AC region, the flow can be modeled using Darcy’s equation with Brinkman correction (Ehlers, 2022)

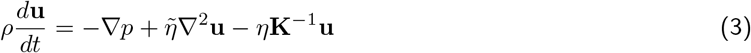

with **K** is the permeability tensor of the AC, and 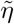 is the effective viscosity in the porous medium. The articular cartilage permeability is dependent on its microstructure. The permeability of the AC region with isotropic collagen arrangement is of the form **K** = *k***I**, with *k >* 0. For the anisotropic region, however, it is not necessarily diagonal. To link microstructure with the permeability, we define the structure tensor (Gasser et al., 2006)

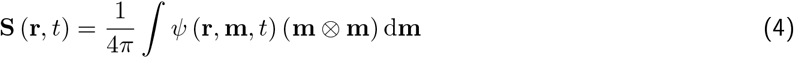

where **m** is the unit vector in an arbitrary direction, *ψ* (**r, m**, *t*) is the orientation distribution function of collagen fibers at spatial location **r** at time *t*, and ⊗ is the dyadic product between two vectors. For an isotropic fiber arrangement, we get **S** (**r**, *t*) = **I**, where **I** is the identity tensor, and for a perfectly aligned fiber arrangement in direction **m**_0_, we get **S** (**r**, *t*) = **m**_0_ ⊗ **m**_0_. In the further section, we will use this structure tensor to differentiate the layers in articular cartilage (section 3.1). Using this description of the articular cartilage microstructure, we consider the following phenomenological dependence of **K** on collagen orientational anisotropy

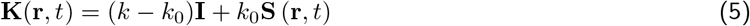

where the first contribution on the right-hand side is due to the isotropic matrix phase, and the second contribution is due to the collagen fiber arrangement. It has to be noted here that in this description, we have not considered any density inhomogeneity in the collagen fiber distribution.

To complete the model description, we also need to specify the boundary conditions. The flexion and extension movements in the joint result in the shear flow between the articulating surfaces. In general, between the two surfaces, there is a lubricating layer of synovial fluid. Given the symmetry in the system, we consider one-half of the joint and model shear flow to be induced by the movement of the plane of symmetry. Therefore, at the plane of symmetry, we have tangential velocity prescribed as one of the boundary conditions (the exact expression to be specified later). Additionally, we have two interfaces - one (at *y* = *h*), between the free space synovial fluid and the superficial layer of the articular cartilage, and two (at *y* = 0), between the articular cartilage and the bone (refer enlarge section of Fig. 1A). We also consider bone to be a porous medium, albeit with different structural and mechanical properties than AC. For simplicity, we assume continuity of both velocity and stress fields at the interfaces, which gives

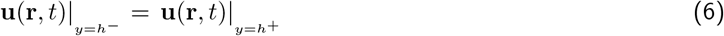

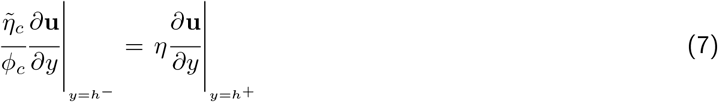

for the interface between the AC and fluid region, and

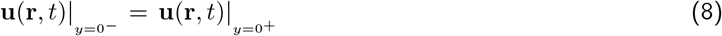

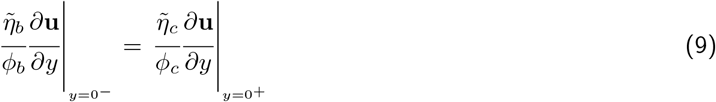

for the interface between AC and bone (Ahmadi et al., 2017) where 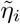 and *ϕ*_*i*_ for *i* ∈ {*b, c*} represent the effective viscosity and porosity of bone and articular cartilage media, respectively.

It has to be noted that the exact nature of the interfacial boundary conditions between fluid-porous media and porous-porous media is an active area of research (Nield, 2009; Ahmadi et al., 2017). Here, we have considered simple forms of interfacial conditions proposed in porous media literature, and for a more detailed analysis, these can be modified to take any specific structural or transport aspect into consideration.

#### Joint movements during embryonic development and post-natal stages

We consider the movement of the joints to be the primary source of mechanical stimulation of the articular cartilage during both embryonic and post-natal stages of development. Given the nature of the environment during the two stages, the movement of the joints during embryonic development and post-natal stages is very different (Pitsillides, 2006; Celik et al., 2023). In development, the joint movements during the early stages are reflexive and sporadic (Celik et al., 2023). They evolve and become more coordinated with the neuromuscular control development (Sharp et al., 1999). The exact nature of these joint movements is simple flexion and extension, especially in limb joints (Rodriguez and Munasinghe, 2016). Therefore, in the model, we incorporate flexion and extension movement of the joint in the form of the following boundary condition

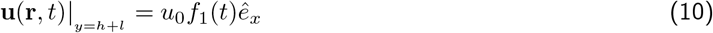

where *u*_0_ is the amplitude of the flexion and extension joint movements and *f*_1_(*t*) is a time-periodic function capturing the exact dynamics of the movement.

In postnatal stages, joint flexion and extension movements are coupled with the normal loading experienced when the animal assumes a standing posture. The effect of the normal loading on the joint will be considered in the following form

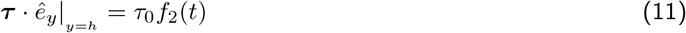

where *τ*_0_ is the amplitude of the pressure on the articular cartilage, and *f*_2_(*t*) is a time-periodic function. This application of pressure on the articulating surface results in compressive deformation of AC along with fluid flow through it. For simplicity, we consider a linear elastic response of the AC matrix during its compression. It has to be noted that, in general, *f*_1_(*t*) and *f*_2_(*t*) may not necessarily be time-periodic as the joint movement may have a much more complex dependence on time. But here, for simplicity, we are considering a periodic dependence of shear and normal loads. Taking inspiration from experimental measurements (top panel in Fig. 1B), we consider alternating compression and shear loading (bottom panel in Fig. 1B) of the joint. The durations and amplitudes *T*_*n*_, *T*_*s*_, and *ϵ*_max_, *γ*_max_ are taken as the model parameters that can be varied to realize different physical activities. It is also worth pointing out here that the normal loading condition, as described in equation (11), is not necessarily the same for all joints. For example, the forelimbs and hindlimb articular joints in the case of kangaroo and humans are expected to experience different normal loadings. We will explore this aspect later in section 3.4.

<u>Now we describe the central hypothesis of this work - the fluid flows generated by the joint movements influence the dynamics of collagen fibers in terms of fiber deposition and removal, with fibers preferentially deposited along the direction of local fluid flow.</u> To model this fluid flow-driven collagen dynamics, we follow two approaches.

#### Collagen fiber dynamics: one-dimensional fiber orientation field model

In the first approach, we model collagen dynamics by describing fiber orientation using a density function *ψ*(**r, m**, *t*)representing the local collagen density (at spatial location **r** and time *t*) of fibers that are oriented along direction **m**. For simplicity, we do not consider any variation in *ψ* along *x* and *z* directions which results in a one-dimensional description of this function *ψ*(*r*, **m**, *t*) where *r* is the depth of articular cartilage. This one-dimensional description can indeed be extended to a more general three-dimensional one, but we are not considering it here.

With this representation of collagen fibers in AC, we assume there is a steady release of collagen protein by the chondrocytes in the articular cartilage. To balance the synthesis of collagen fibers, we also assume their degradation, which follows a first-order kinetics. This assumption is inspired by the experimental observations linking collagen turnover, and not remodeling, with the evolution of collagen structure during post-natal development of articular cartilage (Hunziker et al., 2007). To connect the collagen fiber growth with the synovial fluid flow, we consider that the collagen fiber growth follows the direction of local fluid flow. Therefore, we can write the evolution of collagen fiber density in articular cartilage as

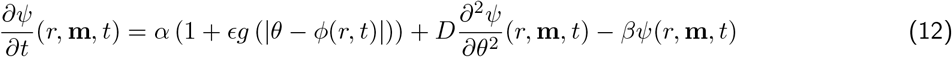

where *ϕ*(*r, t*) is the direction of local fluid flow velocity and *θ* is the orientation of collagen fibers. The first term on the right-hand side of the equation represents the growth of the collagen fibers. For fiber growth along the local fluid flow velocity, an example growth function is the Dirac-delta function *g*(*x*) = *δ*(*x*) that represents fiber growth without any angular dispersion. Given the microscopic scale of collagen fiber assembly, the effect of angular dispersion cannot be ignored. We can imagine angular dispersion to be a combined effect of flow-driven fiber alignment and stochastic noise-induced randomness in the fiber assembly. Therefore, we assume that the angular dispersion in collagen fiber assembly is higher when the flow velocity magnitude is small and *g*(*x*) follows von Mises distribution

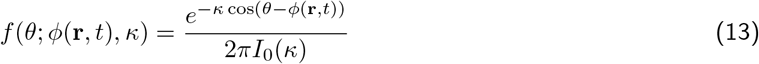

where *κ*, the inverse of orientation dispersion in fibers, depends on the velocity magnitude, and *I*_0_(*κ*) is the modified Bessel function of the first kind of order zero.The second term on the right-hand side of the equation(12) corresponds to the diffusion in the fiber orientations, and the last term corresponds to the fiber degradation. The interplay of these three mechanisms results in a steady state distribution of *ψ*(*r*, **m**, *t* → ∞).

#### Collagen fiber dynamics: three-dimensional discrete fiber model

In the second approach, we deploy a three-dimensional discrete fiber model for the collagen network. We virtually deposit collagen fibers (in the form of straight line segments of length *𝓁*) in a three-dimensional region of AC. The orientation of each line segment follows local synovial fluid flow velocity with some orientational dispersion, as described in the previous section on the one-dimensional model. Here too, each fiber is also degraded at a rate *β* following first-order kinetics. The detailed methodology of this computation model is described in Appendix A.

#### Model analysis and simulation setup

As described above, we follow two approaches to study this system - (i) numerically solving equations describing fluid flow (equations (1)-(3)) and collagen fiber dynamics (equation (12)), and (ii) stochastic simulations of collagen fiber deposition using a discrete fiber model. Fig. 1C shows the steps involved in the simulation of collagen fiber dynamics in both approaches.

In the first approach of solving the equations numerically, we consider the problems of fluid flow and fiber dynamics in an iterative manner (Fig. 1C). We initialize the system with isotropic permeability and estimate the fluid flow. Once the fluid flow is identified, we evolve the fiber dynamics and recalculate the permeability tensor, which now depends on the orientational distribution of the deposited fibers (equation (5)). With the new permeability (anisotropic and inhomogeneous), we recalculate the fluid flow profile and repeat these steps until the solution converges.

In the discrete fiber model, we consider growing collagen fibers as discrete slender filaments. Here, too, we start with isotropic permeability and calculate the fluid flow velocities in different regions of the articular cartilage. For the calculated velocity profile, we grow collagen fibers in all the zones of AC along the local fluid flow direction. At regular intervals, we update the permeability tensor using collagen structural information from the discrete fiber deposition model. We repeat this iterative process to reach convergence.

## 3 Results

### 3.1 Emergence of Benninghoff “arcade” in articular cartilage

Articular cartilage is known to exhibit a highly organized collagen fiber architecture, with a layered structural arrangement known as the Benninghoff “arcade”, first described by anatomist Alfred Benninghoff (Benninghoff, 1925). This structural arrangement, characterized by arc-like collagen fibers that are aligned parallel (*θ* = 0 in Fig. 2A) to the articulating surface in the superficial zone, and oriented perpendicular (*θ* = *π/*2 in Fig. 2A) to the cartilge-bone interface in the deep zone, is believed to contribute significantly to load distribution and joint lubrication (see section 3.4). This three-layered collagen arrangement has been observed in the articulating joints of several organisms, including humans (Fig. 2A), suggesting it plays a fundamental role in cartilage function.

**Figure 2:**
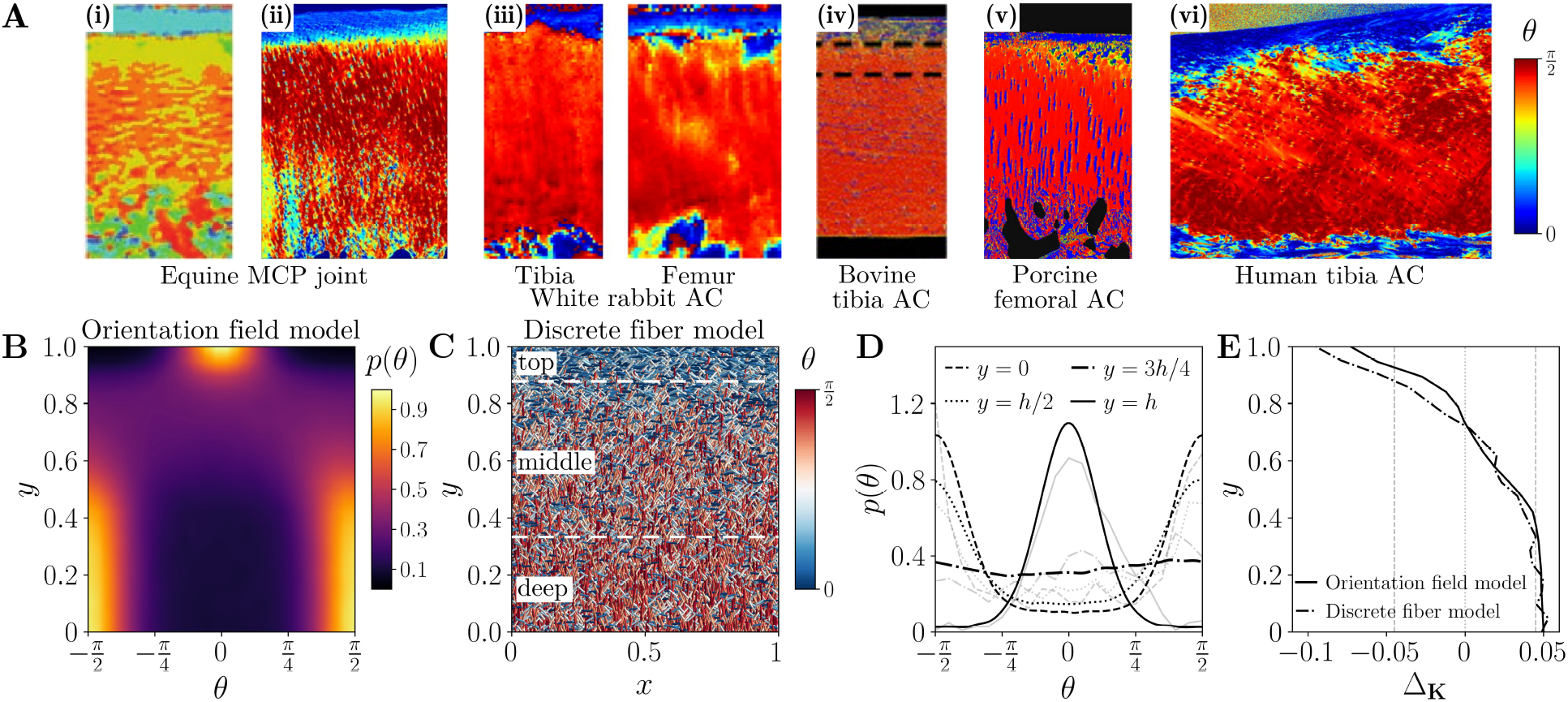
Benninghoff “arcade” architecture of collagen fibers in articular cartilage. (A) shows the maps of the mean orientation angle of collagen fibers in articular cartilage of joints in different organisms - (i-ii) equine AC (images taken with permission from (Brama et al., 2009) and (Oinas et al., 2018), respectively), (iii) white rabbit AC (image taken with permission from (Julkunen et al., 2010)), (iv) bovine cartilage (image taken with permission from (Julkunen et al., 2007)), (v) porcine AC (image taken with permission from (Rieppo et al., 2009)), and (vi) human AC (image taken with permission from (Ebrahimi et al., 2020)). (B) shows the steady state orientation distribution of collagen fibers from the orientation field model. (C) A representative example of collagen network architecture in AC as obtained from the discrete fiber model. (D) The orientation distribution density of collagen fibers at different depths of AC. The dark and light colored lines represent values from the orientation field model and the discrete fiber model, respectively. (E) shows the depedence of permeability anisotropy ratio Δ_**K**_ on AC depth. The vertical dashed lines denote the critical values of Δ_**K**_ marking the boundaries of different layers in AC. For panels (B)-(E), we have *T*_*s*_ = 0.4, *T*_*n*_ = 0.6, *γ*_max_ = 0.25, and *ϵ*_max_ = 0.02.

First, we investigated how mechanical stimuli might give rise to this conserved structure using the two afore-mentioned modeling frameworks: a continuum orientation field model and a discrete fiber-based model. As shown in Fig. 2B, under a combination of normal and shear loading of the joint (simulating a normal walking mode of joint motion), the orientation field model predicts a depthwise evolution of fiber alignment, forming a tripartite architecture analogous to the experimental observations. The superficial zone shows a tangential orientation, the middle zone is more isotropic, and the deep zone features perpendicular alignment relative to the subchondral bone. This observation from the orientation field model is further supported by the discrete fiber model showing the distinct three-layered arrangement of collagen fibers (Fig. 2C). To assess this more quantitatively, we plotted the orientation distribution function at different depths of AC from both modeling approaches. As shown in Fig.2D, at deep regions of AC (*y* = 0 and 1*/*2), the fiber orientation distribution shows a bimodal nature with peaks at *θ* = ±*π/*2. Near the superficial region at *y* = 3*/*4, the distribution of the collagen fiber orientation is uniform, and at the articulating surface *y* = 1, it shows a peak at *θ* = 0, indicating tangential arrangement of the fibers. It should be noted that even though the nature of the collagen orientation is depth-dependent, we did not observe any clear demarcation between different layers. Therefore, to identify the boundaries between the three regions, we calculated the anisotropy ratio

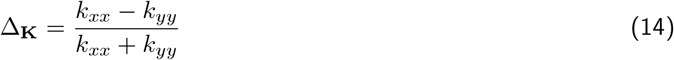

as the ratio of the difference and the sum of directional permeabilities along tangential and normal directions. The dependence of this anisotropy ratio on AC depth, as calculated from both models, is shown in Fig. 2E. We use a critical value of |Δ_**K**_| = 0.045 as the boundaries between different zones, and these are marked in Fig.2E. The choice of the critical value of |Δ_**K**_| is a fitting parameter to ensure anatomically agreeable values of the thicknesses of the three layers of AC. This gives us, for the case reported in Fig. 2B-E, the thickness of the superficial top layer to be approximately 12%, the middle layer to be 54%, and the deep layer to be 33% which is within the reported range of these values (10 ™ 20%, 40 ™ 60%, and 30% respectively (Sophia Fox et al., 2009)).

These findings suggest that depth-dependent fiber alignment in articular cartilage can emerge naturally from local fluid flows generated due to mechanical loading of the joints, as captured by both continuum and discrete modeling approaches. The permeability anisotropy ratio serves not only as a structural marker but also as a potential functional indicator for distinguishing zones with distinct mechanical and transport properties. The mechanical loading necessary to establish the Benninghoff architecture corresponds to the forces experienced during normal adult walking (see section 3.4 for a detailed analysis), which are absent during embryonic development and early post-natal stages. Therefore, we next examine how the collagen structure evolves during these early developmental phases.

### 3.2 Collagen architecture during embryonic development

The role of physical embryonic movement in the development of articular joints is well known (Drachman and Sokoloff, 1966). Particularly, dynamic mechanical stimulation has been shown to be essential for joint cavity formation and cartilage development during embryonic limb growth (Osborne et al., 2002). However, it is important to recognize that the nature of joint loading during development differs from that postnatally. After birth, normal joint loading is primarily driven by body weight during standing and locomotion. In contrast, such weight-bearing forces are minimal during embryogenesis. As a result, internal forces due to limb movements play a more dominant role. We therefore expect shear loading, generated by joint motion, to contribute more significantly to joint development than normal compressive loading.

We investigated the impact of reduced normal loading on collagen fiber organization in articular cartilage to simulate mechanical conditions during embryonic joint development. Figure 3 presents the collagen architecture under two levels of normal load. Under minimal normal loading (*γ*_max_ = 0.2 and *ϵ*_max_ = 0), collagen fibers remain predominantly aligned parallel to the articular surface in the superficial zone and become more isotropic with depth (Fig. 3A–B). This pattern is reflected in the depthwise permeability anisotropy, which remains close to zero (inset in Fig. 3B). These results align with experimental observations in neonatal sheep, where collagen fibers are oriented tangentially at the surface (left panel in Fig. 3C, see (van Turnhout et al., 2010)). Given the idealized representation of AC, model’s predicted mean fiber orientation (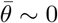 at all AC depths) is also in agreement with experimental data which shows 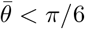 (right panel in Fig. 3C).

**Figure 3:**
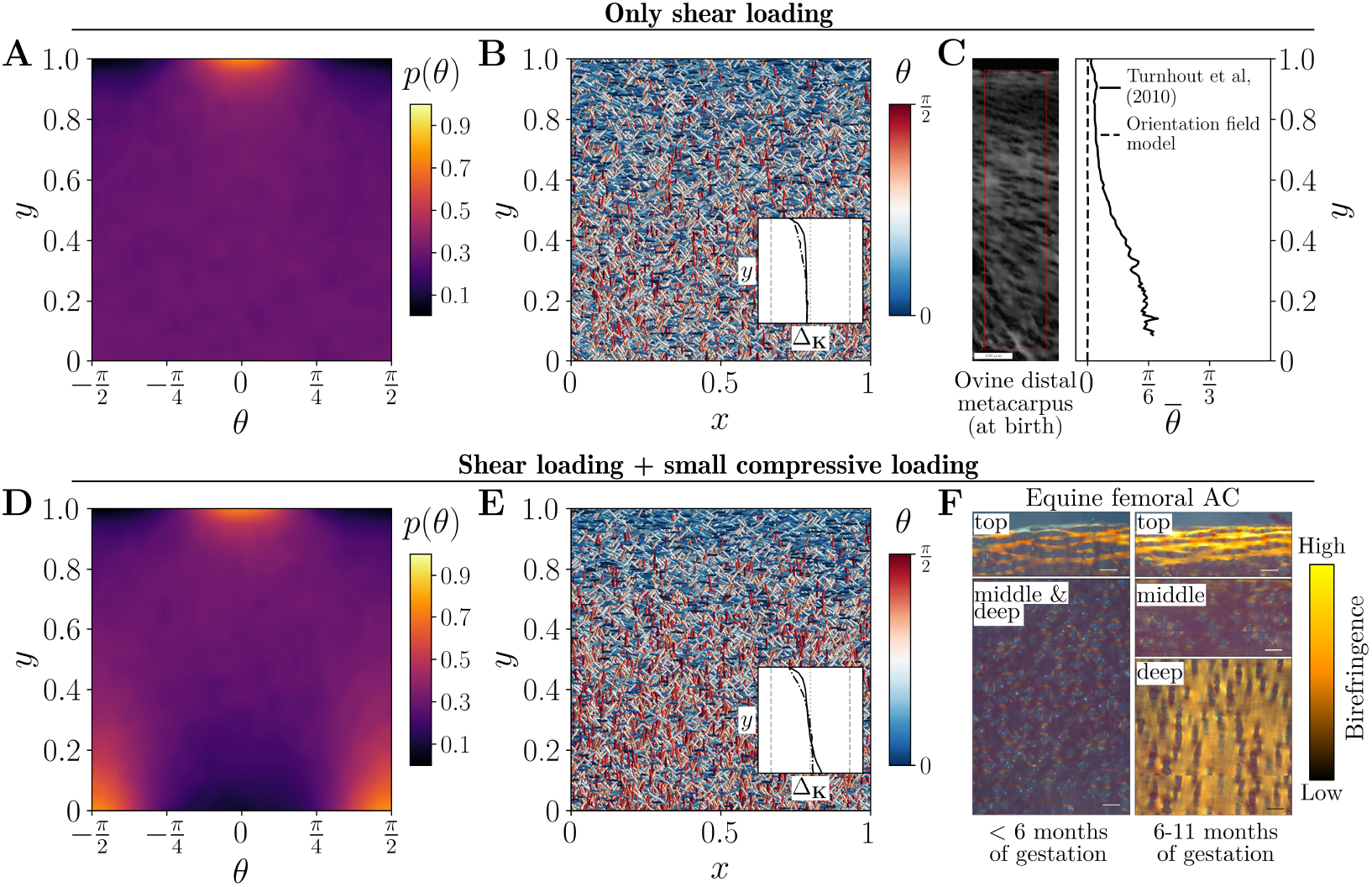
Collagen fiber arrangement in articular cartilage during embryonic development. The steady-state orientation distribution of collagen fibers, as predicted by the orientation field model (A) in the absence of compressive loading in the joint, (D) when a small amount of compression is applied along with shear on the developing joint. Representative plots showing collagen fibers for these two cases, as obtained from the discrete fiber model, are shown in (B) and (E), respectively. Corresponding insets show the dependence of permeability anisotropy ratio Δ_**K**_ on AC depth. (C) Experimental measurement of collagen fiber orientation in neonatal ovine distal metacarpus (image taken with permission from (van Turnhout et al., 2010)). The left panel shows the polarised light microscopy image and the right panel presents the quantification of average collagen fiber orientation. (F) Experimental images showing birefringence pattern of collagen fiber orientation at two gestation stages for equine femoral AC (images taken with permission from (Cluzel et al., 2013)). For panels (A)-(C), we have *T*_*s*_ = 0.6, *T*_*n*_ = 0.15, *γ*_max_ = 0.2, and *ϵ*_max_ = 0. For panels (D)-(E), we have set *ϵ*_max_ = 0.002 while keeping all other parameters to be same as those in (A).

With a small increase in normal loading (*ϵ*_max_ = 0.002), collagen fibers begin to align perpendicular to the articular surface in the deep zone, indicating the emergence of a three-layered architecture in articular cartilage (Fig. 3D–E). However, the degree of alignment in each zone remains lower than that observed in mature tissue, as reflected by the lower Δ_**K**_ values compared to adult cartilage (compare inset of Fig. 3E and Fig. 2E). This trend is consistent with experimental observations from neonatal horse knee joints (Fig. 3F, see (Cluzel et al., 2013)). In early gestation (< 6 months), no clear zonal architecture is observed (left panel), whereas by approximately 11 months of gestation, collagen fibers exhibit a well-organized, three-layered structure (right panel). It has to be noted, however, that despite a layered arrangement, the degree of fiber alignment in the top and deep layers of AC remains weaker than that in adult tissue (more details in the next section). These observations suggest that the evolution of collagen architecture during embryonic development can be attributed to changing mechanical loading conditions on joints over the course of development.

### 3.3 Post-natal evolution of collagen architecture in different joint types

Following birth, the mechanical environment of joints changes markedly due to weight-bearing and increased locomotor activity. These post-natal loading conditions can play a critical role in further organizing the collagen architecture of AC. Therefore, we explored the evolution of collagen arrangement in two joint types - first, joints with high compressive and shear loading (for example, knee joint), and second, joints with a low degree of compressive loads (such as elbow joints) to highlight how joint-specific loading patterns influence post-natal AC architecture. Fig. 4 shows the time course evolution of collagen arrangement in AC of joints with high compressive loading as obtained from the models and experiments. As shown in the previous section, at birth, the collagen arrangement is weakly aligned along the articulating surface. It evolves to attain the three-layered Benninghoff architecture on maturity (Fig. 4A). This evolution is also reflected in the increasing density of fibers aligned normal to the articulating surface (*θ* = *π/*2) as seen in Fig. 4B. Correspondingly, there is a progressive reduction in the thickness of the superficial collagen layer with fibers oriented parallel to the surface (*θ* = 0), indicating a rearrangement of fiber orientations during post-natal growth.

**Figure 4:**
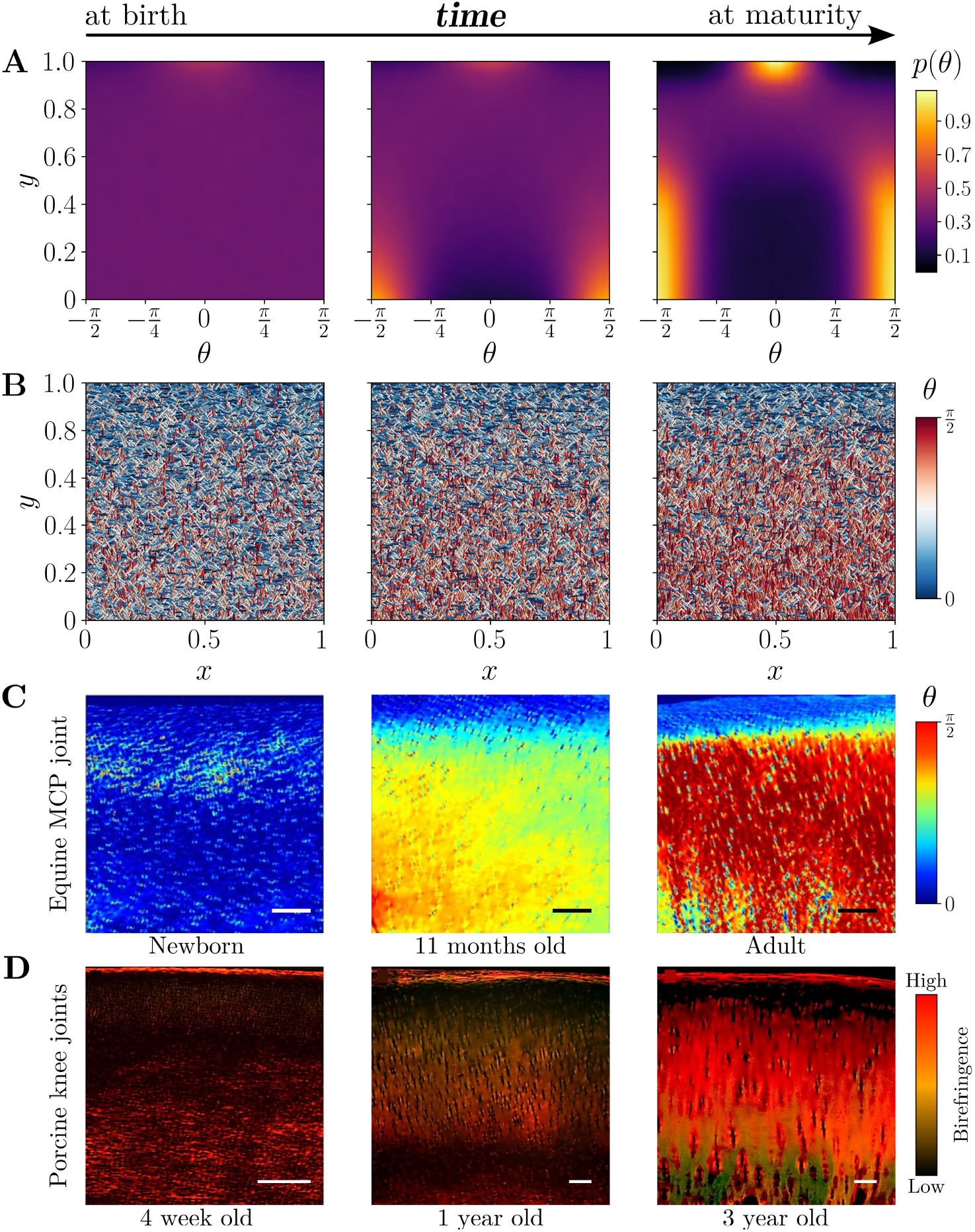
Post-natal evolution of collagen architecture in articular cartilage. (A) shows the evolution of the steady-state orientation distributions of collagen fibers at three distinct post-natal developmental stages. The corresponding arrangements of collagen fibers obtained from the discrete fiber model are shown in (B). Please note that the first and last panels in (A) and (B) are a repetition of Fig. 3A-B and Fig. 2B-C, respectively, and are given here for comparison. (C) Experimental images showing the mean orientation angles of collagen fibers in equine articular cartilage during maturation, obtained via polarized light microscopy (reproduced with permission from (Oinas et al., 2018)). (D) shows experimental images of the birefringence pattern, highlighting regions of low (dark) and high (bright red) collagen alignment at various developmental stages of porcine knee articular cartilage (reproduced with permission from (Gannon et al., 2015)). All scale bars in the experimental images indicate 100 µm. The parameter values for simulations for AC at birth and at maturity are set as those in Fig. 2B and Fig. 3A, respectively. For the intermediate state, we have *T*_*s*_ = 0.5, *T*_*n*_ = 0.4, *γ*_max_ = 0.25, and *ϵ*_max_ = 0.006.

Experimental evidence for this transition is shown in Fig. 4C-D, which shows mean fiber orientation angles (as obtained from polarized light microscopy) and fiber alignment (as obtained from birefringence) for equine metacarpophalangeal joints and porcine knee joints, respectively. Both of these cases show a clear post-natal shift in collagen fiber arrangement - starting from fiber weakly aligned along the articulating surface (first panels in both rows), we observe the establishment of Benninghoff architecture in both joints at the maturity of the animal (last panels in both rows). It has to be noted that the mean fiber orientation plot (as shown in Fig. 4C) does not depict the magnitude of dispersion in fiber orientation. However, in the birefringence plot shown in Fig. 4D, the zonal architecture of collagen fiber arrangement is clearly visible for adult animals.

In contrast to the knee or femoral joints that are under constant compressive loading post-natally, elbow joints in bipedal animals (for example, kangaroos and humans) do not experience that magnitude of compressive stresses. Therefore, in these joints, the fluid flow is expected to have a lesser contribution in the direction normal to the articulating surface, and this should be reflected in the collagen arrangement in the AC of the elbow joints. We performed numerical simulations using orientation field and discrete fiber models with reduced mechanical loading (relative to a knee joint), particularly in the normal direction, to mimic the mechanical stimulation of an elbow joint in bipedals. Fig. 5A-B show the steady state fiber orientation distribution and a representative example of fiber arrangement for an elbow joint, respectively. Given a lower magnitude of the mechanical load, the dispersion in the orientation distribution is higher in this case compared to AC in the adult knee joint (comparing Fig. 2B with Fig. 5A). This is also seen in experimental observations on AC in the elbow joints in kangaroos, as shown in Fig. 5C. A comparison of collagen fiber orientation in the AC of the femoral condyle and distal humerus reveals distinct patterns - the femoral condyle exhibits prominent fiber alignment in both the top (dark green) and deep zones (dark red), whereas the distal humerus shows fiber alignment, but with notably lower degrees of organization in these two regions (light green and light red, respectively).

**Figure 5:**
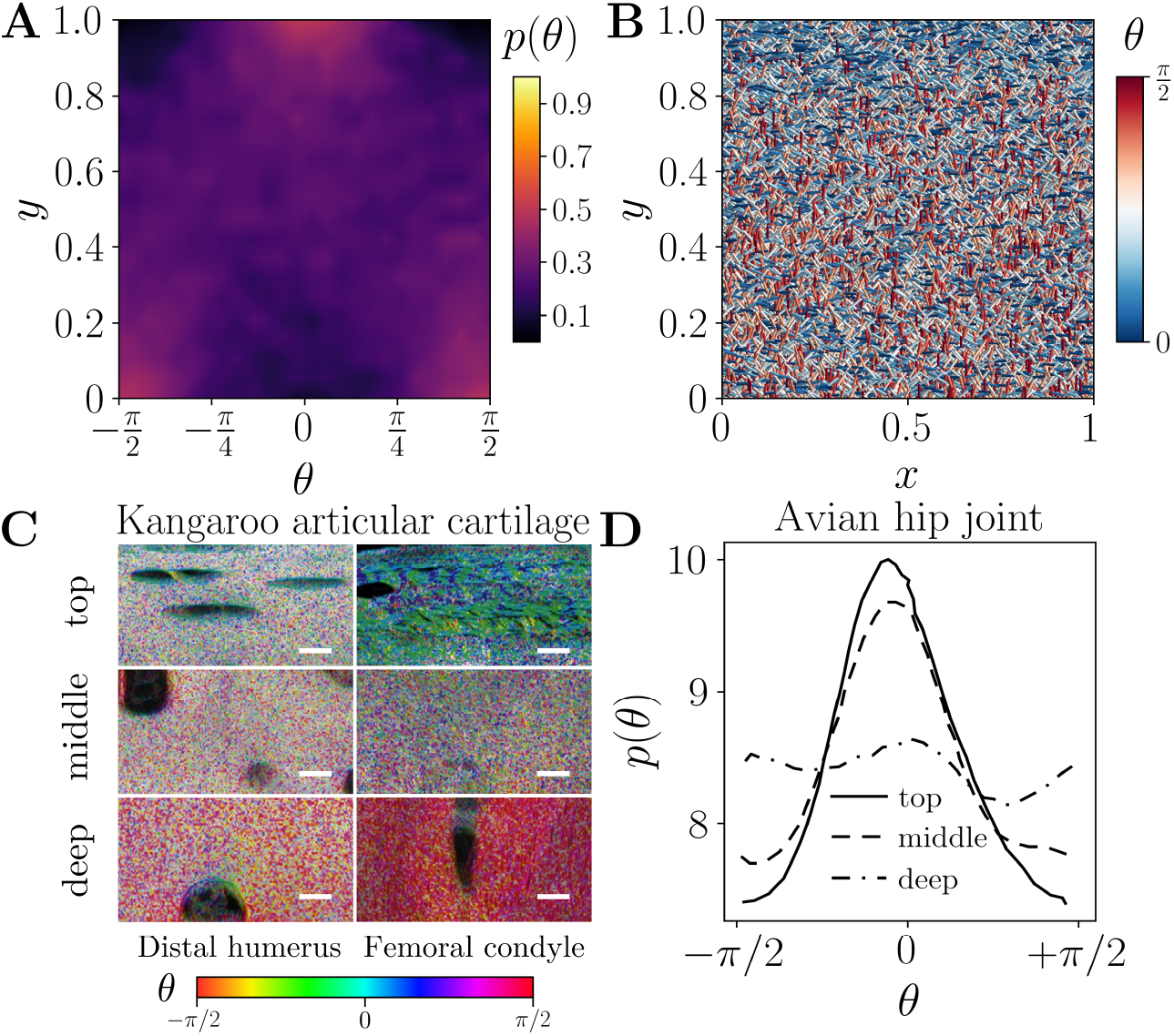
Post-natal collagen architecture in elbow joints. (A) shows the steady-state orientation distributions of collagen fibers in elbow joints, and the corresponding collagen network obtained from the discrete fiber model is shown in (B). (C) Experimental images showing orientation of collagen fibers in the articular cartilage of the distal humerus and the femoral condyle of adult kangaroos (reproduced with permission from (He et al., 2013)). (D) shows the deconvolved profiles of the fiber distribution in avian cartilage at different depths (data taken from (Wess et al., 1997)). All scale bars in panel (C) indicate 10 µm. For (A) and (B), we have *T*_*s*_ = 0.4, *T*_*n*_ = 0.1, *γ*_max_ = 0.08, and *ϵ*_max_ = 0.001.

Furthermore, experimental explorations of articular cartilage in avian hip joints have reported a collagen fiber architecture different from Benninghoff’s arcade pattern (Wess et al., 1997). As shown in Fig. 5D, in the turkey antitrochanter, the collagen fiber orientation in the top and middle zones is predominantly aligned parallel to the articulating surface. This can also be attributed to the lesser extent of the compressive loading, given the anatomical structure of the avian hip joint.

These findings collectively demonstrate that post-natal mechanical loading plays a decisive role in shaping the collagen architecture of articular cartilage. While high compressive and shear loading, as seen in knee joints, promotes the development of the classical Benninghoff structure, joints subjected to lower compressive stresses, such as the elbow in bipedals or avian hip, exhibit more dispersed or surface-parallel fiber arrangements. This highlights the critical interplay between joint-specific biomechanics and tissue microstructure during post-natal development.

### 3.4 Effect of physical activity on collagen architecture

As highlighted earlier, physical activity has been known to influence the articular cartilage structure (Vincent and Wann, 2019). In fact, a consistent regimen of joint loading during youth has been shown to significantly reduce the risk of developing osteoarthritis (Helminen et al., 2000). Therefore, with the present modeling framework, we studied the effect of the nature of physical activities on collagen architecture in articular cartilage.

We explored different physical activity regimens by varying the durations (*T*_*s*_ and *T*_*n*_) of the applied shear and compressive loads on the joints. Of course, the exact nature of any physical activity involves more complexity in terms of the magnitude of the compressive and shear forces. But, for simplicity, we have considered the duration of the mechanical loading to be the representative of that particular physical activity. This helps in keeping the parameter space to be two-dimensional, which can be explored systematically. In principle, one can extend this approach by incorporating different combinations of loading magnitudes and durations (in a four-dimensional phase space). Using this setup, we performed numerical simulations with different durations of compressive loading *T*_*n*_, and shear loading *T*_*s*_ to the AC and calculated the thicknesses of the three zones in the AC using the approach described earlier (section 3.1).

Fig. 6A shows the thickness of three zones in *T*_*s*_-*T*_*n*_ space. For no activity or an idle joint, the durations of both compressive and shear loading of the joints are very small (case-N in Fig. 6A-B). In this case, we observe that most of the AC has a random fiber arrangement, resulting in a very thick middle zone and negligible thicknesses of the top and deep zones.

**Figure 6:**
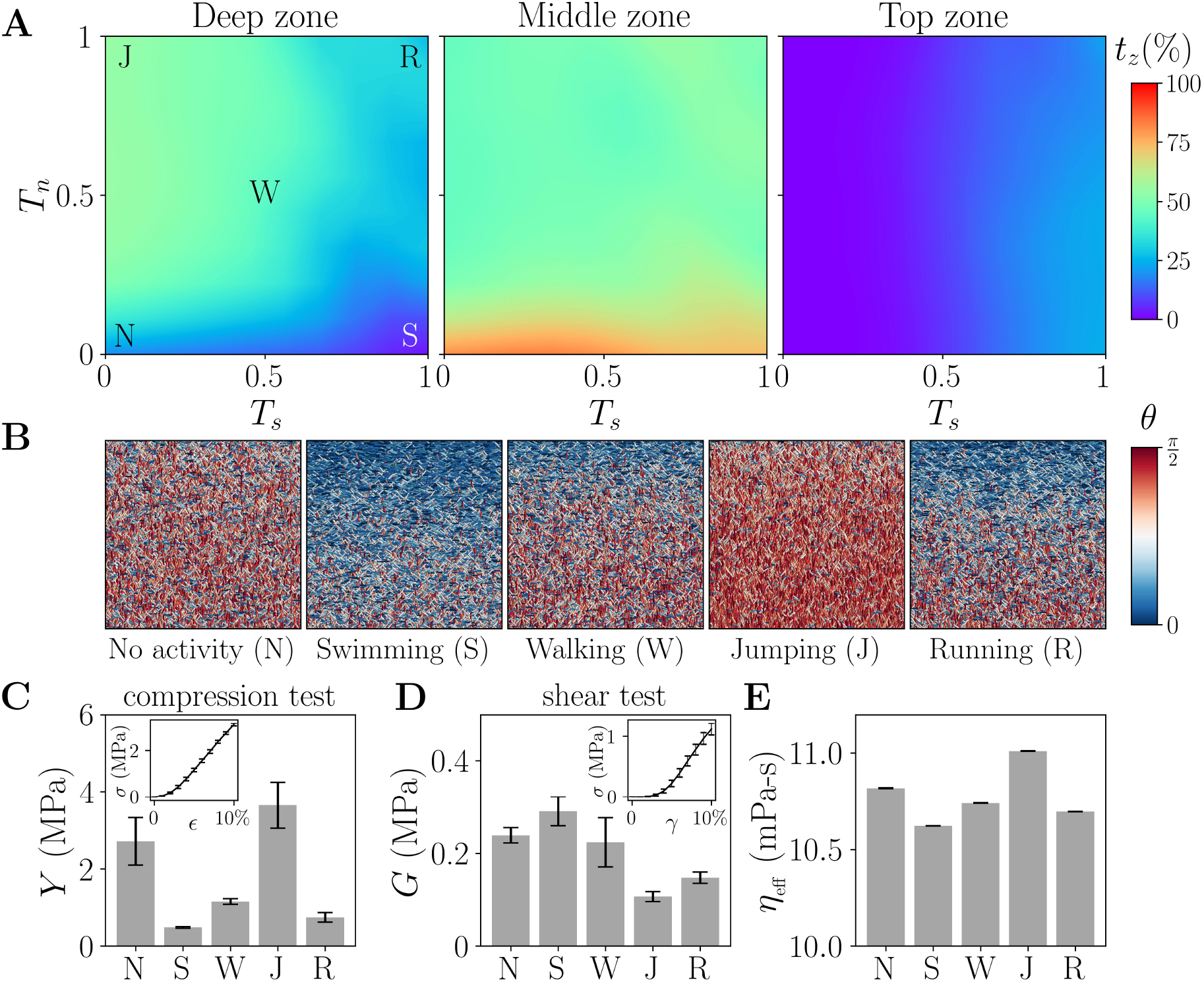
Effect of physical activity on articular cartilage structure and function. (A) shows the thickness of three zones in articular cartilage under varying mechanical loading conditions. The representative collagen networks obtained using the discrete fiber model under five distinct loading conditions, representing - no activity (N), swimming (S), walking (W), jumping (J), and running (R) - are shown in (B). The moduli of representative examples of AC are shown under (C) uniaxial compression and (D) shear deformation. The corresponding insets in (C) and (D) show the stress–strain plot for one of the physical activities (walking). (E) shows the effective viscosity of the synovial fluid through AC under these five representative loading conditions. Here we have fixed the values of *γ*_max_, and *ϵ*_max_ to be same as those in Fig. 2B.

In the second extreme scenario, where compressive loading is negligible (case-S, swimming can be a representative example with this loading), the deep zone disappears and the AC has a relatively thick top and middle zones. Similarly, for no shear loading (case-J, for example, jumping), the top zone is not seen in AC, and it is found to be equally spanned by the middle and deep zones. For equal contributions from shear and compressive loads (cases-W and R, activities like walking and running, respectively), the AC shows a three-layered Benninghoff architecture, albeit with different zonal thicknesses. These observations underscore that the zonal organization of collagen architecture in AC is highly sensitive to the nature and balance of mechanical loading, with distinct activity patterns giving rise to characteristic structural adaptations.

To evaluate the physiological implications of these structural adaptations in AC to physical activities, we estimated its compressive (*Y*) and shear (*G*) moduli, along with the effective viscosity (*η*_eff_) of synovial fluid during joint motion. The elastic moduli quantify load-bearing capacity, while the effective viscosity captures joint lubrication-both essential for normal joint function. The details of the methodology are given in Appendix B and Appendix C. Fig. 6C-E shows these quantities for AC evolved under different physical activity regimens. Both compression and shear deformations of collagen fiber reinforced AC show a nonlinear strain-stiffening response (insets in Fig. 6C-D, also see Fig. S1 in supplementary materials). For articular cartilage subjected to moderate compression and shear (simulating walking, case-W), we observe compressive and shear moduli to be on the order of 1 MPa and 0.1 MPa, respectively. Experimental measurements support these values, with reported ranges of 0.1-10 MPa for compressive modulus (Kabir et al., 2021) and 0.01–1.0 MPa for shear modulus (Krakowski et al., 2024; Wyse Jackson et al., 2022). The wide variability in these moduli arises from differences in joint type, anatomical location, and species. For an AC with no physical activity we observe a stiffer response under compression and shear. Articular cartilage subjected to excessive compressive (case-J) and shear loading (case-S) exhibits contrasting elastic properties. Specifically, case-J shows a high compressive modulus and low shear modulus, whereas case-S displays the opposite trend. These differences can be attributed to the predominant fiber orientations - a greater fraction of fibers oriented normal to the articulating surface in case-J supports compressive loads, while a higher fraction of fibers aligned parallel to the surface in case-S enhances shear resistance. Finally, the AC with high magnitudes of compression and shear loads (case-R) demonstrates marginally lower elastic moduli. Across all cases, the effective synovial fluid viscosity exhibits a trend that appears to correlate with collagen fiber orientation. A higher fraction of fibers aligned parallel (perpendicular) to the articulating surface results in a slightly lower (higher) effective viscosity, although the differences are modest (Fig. 6E). These results suggest that a balanced mechanical environment involving moderate levels of both compression and shear, such as during walking, optimally tunes the structural and rheological properties of AC, enhancing both its load-bearing capacity and lubrication efficiency.

It is important to note that this analysis focuses on the influence of mechanical loading, arising from different physical activities, solely on collagen fiber architecture. In vivo, joint loading also impacts other physiological processes such as nutrient transport, suggesting that the observed outcomes likely result from the combined effects of transport and fiber remodeling. Additionally, we have assumed unchanged chondrocyte physiology under varying loading conditions, though experimental studies indicate that excessive or no mechanical loading can alter cell behavior (Chang et al., 2019). While prior experiments have examined the consequences of joint immobilization (Maldonado et al., 2013; Vanwanseele et al., 2002; Nagai et al., 2015), the differential effects of specific loading modes (e.g., compression vs. shear) on AC structure remain experimentally unexplored. Hence,further experimental validation is necessary to test the model predictions presented here.

### 3.5 Collagen architecture in osteoarthritis

To understand how degenerative changes impact collagen organization, we examined the fiber architecture of articular cartilage in the early stages of osteoarthritis. Specifically, we explored how localized damage and altered mechanical environments influence collagen remodeling and contribute to disease progression. To model the effect of OA, we modified the discrete fiber model to include the effect of damaged AC in the form of an altered fluid flow field. The resultant fluid flow in the AC is given by

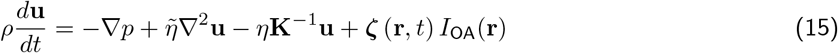

where the last term represents the effect of osteoarthritis-induced disruption on local fluid flow. Here,

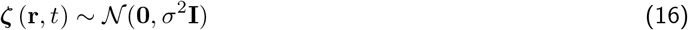

is a vector-valued Gaussian white noise with zero mean and isotropic covariance *σ*^2^**I**, uncorrelated in space and time. The spatial indicator function

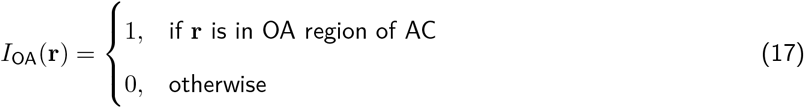

ensures that the stochastic perturbation is localized to OA-affected regions only. Using this framework, we explored the evolution of the collagen fiber arrangement in AC with early and advanced OA.

Fig. 7A shows the collagen fiber organization in AC samples with early and advanced OA. The regions of the AC affected by osteoarthritis are marked with a white dashed line. In both cases, in the OA-affected regions in the top zone, the collagen fibers get reorganized in random orientation as opposed to an aligned arrangement in healthy AC. This is also observed experimentally in OA-affected human tibial articular cartilage where the top zone of AC has isotropic fiber orientation (marked by scattered irregularities in the top zone in Fig. 7C). This can be appreciated by comparing the mean orientation of collagen fibers shown in Fig. 7C to those shown in Fig. 2A or Fig. 4C). The effect of the OA induced randomness in the synovial fluid flow, however, does not remain confined to that region. Instead, the AC surrounding the osteoarthritic region also has its collagen fibers disoriented from their healthy arrangement. This can be seen in Figs. 7B and 7D from the collagen fiber orientation distribution and mean collagen fiber orientations at different AC depths. In early OA, collagen fiber orientation becomes disrupted in the top zone of AC (Fig. 7B), with the disorganization worsening as OA progresses. In fact, with progressing OA, its effect on collagen architecture is also seen in the middle zone as the mean fiber orientation is deviated from that for a healthy AC (Fig. 7D solid line vs other lines).

**Figure 7:**
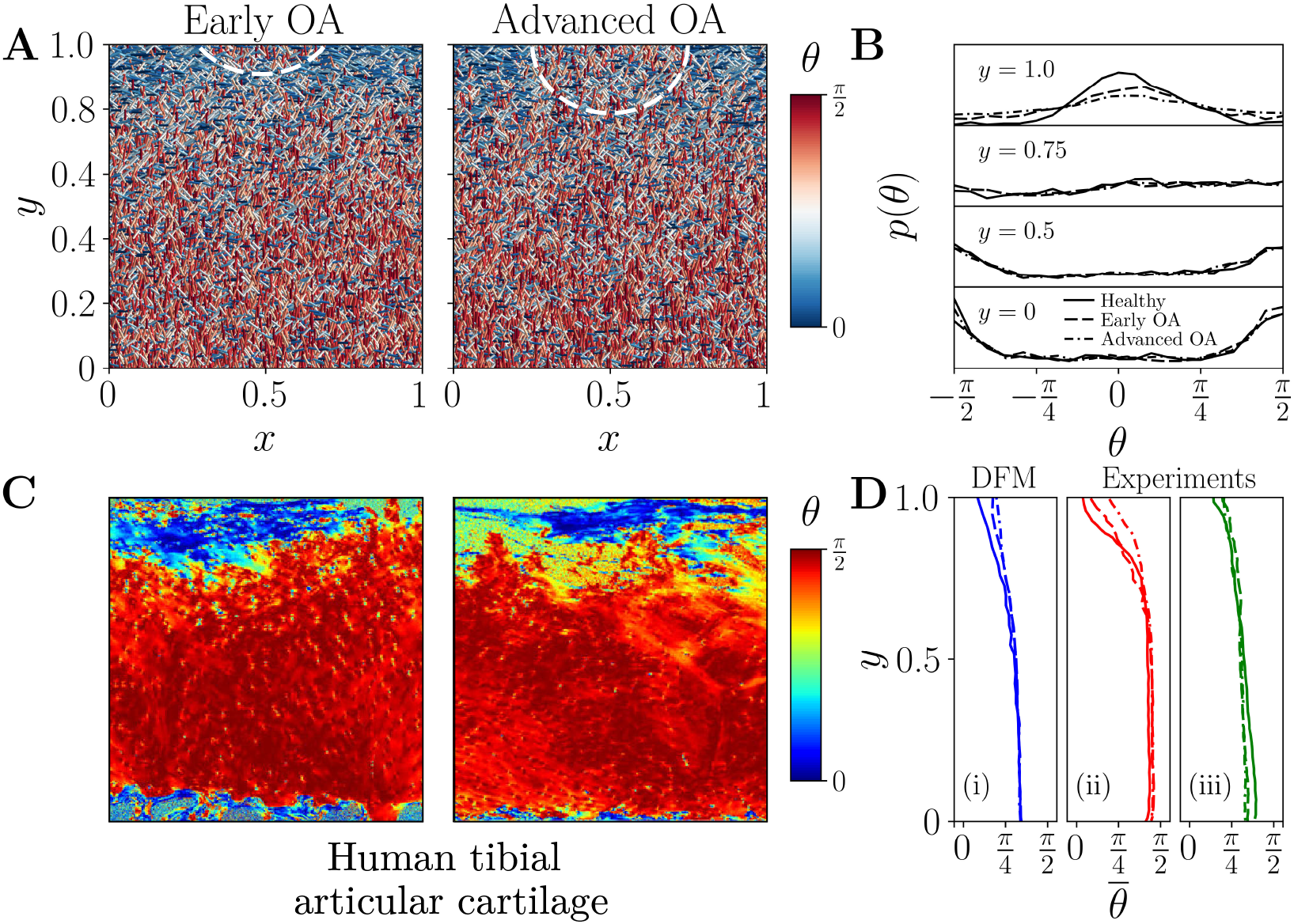
Articular cartilage structure in osteoarthritis. (A) shows the collagen architecture in AC during early OA (left) and advanced OA (right) as obtained from the discrete fiber model. The white dashed lines mark the region affected by OA in both. The orientation distribution density of collagen fibers at different depths of OA-affected AC is shown in (B). (C) Experimental images showing mean collagen orientation angle in early OA (left) and advanced OA (right) cartilage (reproduced with permission from (Ebrahimi et al., 2020)). (D) Comparison of depth-wise collagen mean orientation angle 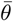 between the (i) discrete fiber model and (ii-iii) experimental observations (data for (ii) and (iii) are taken from (Ebrahimi et al., 2020) and (Saarakkala et al., 2010), respectively). (B) and (D) share the same line types for healthy AC, and AC with early and advanced OA. Here, the mechanical loading parameter values for the simulations are the same as those in Fig. 2B.

These results suggest that the structural disruption caused by OA is not limited to the visibly damaged regions but propagates into the surrounding tissue, where the native collagen architecture might get affected. This progressive disorganization of fiber orientation may impair the mechanical integrity (in terms of the elastic moduli, see Fig. 6C-D) and physiology of AC (in terms of friction between the two surfaces, see Fig. 6E), and contribute to the feed-forward deterioration characteristic of osteoarthritic joints.

## 4 Discussion

Mechanical stimulation is essential for joint development and maintenance (Decker et al., 2015; Decker, 2017; Sun et al., 2011), yet the specific cues driving morphogenesis and long-term homeostasis have remained unclear. Disruption of joint formation under both rigid and flaccid paralysis suggests that dynamic, not static, loading is critical (Brunt et al., 2016). In this work, we show that a plausible mechanism for translating these dynamic forces into microscale tissue organization is synovial fluid flow during joint motion (Fahy et al., 2018). The role of fluid flow in extracellular matrix (ECM) remodeling, particularly in promoting ECM alignment along the direction of flow, has been well recognized in the tissue engineering literature (Lee et al., 2006; Ahmed et al., 2022). Building on this, we extend the concept to developmental and physiological contexts by integrating computational simulations with experimental data and tracing the progression of collagen organization from early embryonic stages through post-natal development.

Our analysis shows that the zonal organization of the Benninghoff collagen fiber architecture arises from the spatial variation in the directions of synovial fluid flow within the articular cartilage. In the superficial zone, flow is predominantly parallel to the articulating surface, while in the deep zone, it is primarily normal to the surface.The middle zone exhibits a transition between these extremes, with time-averaged flow appearing isotropic. This zonally varying flow pattern drives the emergence of the Benninghoff structure in mature joints (Fig. 2). While the zonal arrangement of collagen fibers is widely accepted, it is not universally observed across all joint types or species. For example, weak zonal organization is evident in the kangaroo distal humerus (Fig. 5C), and largely absent in the avian hip joint (Fig. 5D). Our fluid flow–driven model suggests that these variations arise from differences in joint-specific mechanical loading across organisms. Such joint- and organism-specific collagen architectures give rise to articular cartilage that is functionally optimized for mechanical load-bearing and efficient lubrication during joint movement (Fig. 6).

Conflicting reports exist on neonatal AC structure, with some studies indicating isotropic properties (Brama et al., 2000; Brommer et al., 2005; Brama et al., 2002), while others report zonal organization without the characteristic Benninghoff architecture (van Turnhout et al., 2010; Cluzel et al., 2013). These differences likely stem from variations in mechanical stimulation during embryonic development. Compared to post-natal stages, joint loading in the embryo is significantly lower, with spontaneous movements generating shear-dominated flow but minimal compressive loading – as seen in the equine femur (Fig. 3F) or entirely absent in the ovine metacarpus (Fig. 3C). Such weak compression can lead to collagen fibers aligning parallel to the articulating surface, even within the middle zones of AC at birth.

Several studies have examined the impact of mechanical loading, via exercise or joint immobilization, on articular cartilage structure, demonstrating that collagen fiber orientation is influenced by loading in a site-specific manner (Brama et al., 2000, 2002; Brommer et al., 2005; Brama et al., 2009). These effects are generally attributed to local adaptations of AC to sustained mechanical forces (Brama et al., 2009), yet such explanations remain phenomenological and lack mechanistic details that can provide a control over collagen fiber arrangement. A recent study using cartilage organoids identified tissue growth direction as a potential driver of parallel collagen fiber alignment (Peters et al., 2024). However, this study lacked applied mechanical stimulation and does not fully account for the collagen arrangements observed during embryonic development (van Turnhout et al., 2010). Another study using bone-cartilage explants subjected to brief compressive loading reported maximal changes in collagen architecture in the middle zone (Hu et al., 2025). While this appears to contrast with our findings, where the deep zone shows prominent remodeling under sustained oscillatory compression, the difference likely reflects the role of timescales, where the explant study captured short-term passive reorientation of collagen fibers, whereas our work focuses on long-term, load-induced remodeling. In another finite element–based analysis,collagen fiber orientation was predicted based on the directions of principal strains (Wilson et al., 2006). They successfully reproduced the zonal collagen architecture observed in adult articular cartilage. However, their study considered only compressive loading and neglected the contributions of interstitial fluid flow and shear stresses, which are expected to play a critical role in shaping AC structure, particularly in joints undergoing flexion–extension movements.

In early-stage osteoarthritis, even minor cartilage damage can disrupt local fluid dynamics, leading to erratic flow patterns and altered permeability at the injury site. Our findings suggest that such changes trigger a feed-forward remodeling of collagen fibers, compromising the tissue’s ability to bear load effectively. The consistency of these results with experimental observations in OA-affected joints (Fig. 7) points to potential therapeutic opportunities, particularly in the early stages of degeneration. By leveraging the fluid flow–driven mechanism proposed here, optimal mechanical loading regimens may be identified to promote repair and restore functional collagen architecture. For late-stage OA and other degenerative diseases of the AC, one of the therapeutic strategies is tissue engineering (see (Mukherjee et al., 2025; Welsh and Sikder, 2025; Yang et al., 2024) for some recent reviews). While the importance of mechanical stimulation in articular cartilage engineering is well recognized (Fahy et al., 2018), the specific loading regimes required to direct collagen organization and restore function remain poorly defined. The fluid flow–driven mechanism of collagen remodeling proposed in this study offers a framework to identify the directionality and type of mechanical cues essential for promoting zonal architecture and tissue maturation. These insights hold direct relevance for advancing tissue engineering approaches, particularly in the rational design and mechanical conditioning of bioreactors.

While the present model captures key mechanobiological drivers of collagen organization in articular cartilage, several important factors remain to be explored to develop a more comprehensive and physiologically realistic framework. First, the simplified geometry used here does not account for the complex three-dimensional contours of native joints, which influence local mechanical environments. Additionally, the loading conditions applied in this study are idealized representations of activity-induced forces; in reality, joints experience a broad spectrum of dynamic, multi-axial loads that vary in frequency and magnitude. Incorporating the effects of cartilage growth, particularly during development and maturation, would further enhance the model’s ability to capture evolving tissue architecture. Chondrocyte density, which influences matrix production and remodeling, and nutrient transport dynamics, especially in avascular cartilage, also warrant consideration. Moreover, the synovial fluid was modeled as Newtonian, although its non-Newtonian behavior significantly affects joint lubrication under varying shear rates. Finally, in the context of osteoarthritis, incorporating the spatiotemporal progression of collagen damage could provide mechanistic insights into disease initiation and progression.

Mechanical stimulation during embryonic development plays multiple roles beyond influencing collagen organization, including guiding joint morphogenesis and cavitation (Osborne et al., 2002), which can make experimental validation of structure-specific hypotheses particularly challenging during early developmental stages. Furthermore, another aspect that is not considered in the model is the effect of mechanical loading on AC metabolism. For instance, patients with spinal cord injury exhibit elevated levels of cartilage degradation markers (Vanwanseele et al., 2004; Findikoglu et al., 2012), and animal studies have consistently demonstrated that lack of mechanical stimulation leads to progressive cartilage degeneration (Tomiya et al., 2009). The findings of our study further suggest that, in addition to preserving cartilage integrity, mechanical stimulation modulates the evolving architecture of the tissue over time. Collectively, these insights highlight the central role of joint mechanics in both the development and long-term function of articular cartilage.

## 5 Conclusions

In this work, we have presented a fluid flow–driven mechanism for the evolution of collagen architecture in articular cartilage, integrating developmental biomechanics with fiber remodeling. We have shown that during the late embryonic and postnatal stages, joint motion–induced shear and compressive loading differentially regulate collagen orientation and zonal organization. Notably, moderate levels of both shear and compression, characteristic of walking, yield AC with balanced mechanical properties, that is, higher load-bearing capacity and efficient lubrication. The model also replicated the structural features observed across different species and joint types and predicted progressive collagen disorganization under osteoarthritic conditions, providing a mechanistic basis for early degeneration. These insights not only present a biomechanical mechanism for the emergence of characteristic Benninghoff architecture of collagen fibers in AC but also inform tissue engineering efforts by identifying mechanical environments that are conducive for the regeneration of AC.

## Acknowledgements

MSR gratefully acknowledges the late Dr. Anupam Pal for his insightful discussions, which were instrumental in shaping the ideas presented in this work. Financial support from the Science and Engineering Research Board (Startup Research Grant SRG/2021/001020), Government of India, is also acknowledged. DJM acknowledges financial support from the Ministry of Education, Government of India.

## Appendix A Discrete fiber model

This appendix outlines the framework for the numerical model to generate articular cartilage using the discrete fiber model (DFM) of collagen fibers in response to synovial fluid flow in the background. The model incorporates stochastic fiber deposition and removal from the domain until a steady-state is reached. In the numerical simulation of collagen fiber network generation at each time step, some fibers are added and some existing fibers are removed. The number of collagen fibers added to the network, *α* is taken from a Gaussian distribution, that is

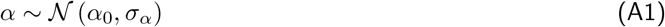

where *α*_0_ represents the average number of fibers deposited at each time step and *σ*_*α*_ is the standard deviation of the distribution. The parameters *α*_0_ and *σ*_*α*_ can be chosen to reflect biological factors such as chondrocyte activity and/or pathological conditions. At each iteration *α* (rounded to the nearest integer), seed points are uniformly generated in the three-dimensional domain of size *L* × *W* × *H*, representing a small section of AC (Fig. A1A). For each seed location **r**_0_(*x, y, z*), a straight fiber of length *𝓁* is added to the domain at an orientation *θ*_*f*_ (**r**, *t*) (A1B). As mentioned earlier, the collagen fiber deposition follows the local fluid flow velocity. Therefore, the fiber orientation *θ*_*f*_ depends on the local fluid flow direction at that instant at time *t*, that is *ϕ*(**r**_0_, *t*), which is obtained from the velocity of the fluid in the articular cartilage using equation (3). As described earlier, given the microscopic nature of the process, we also consider a finite amount of dispersion and *θ*_*f*_ is taken from von Mises distribution (equation 13) centered at *ϕ*(**r**_0_, *t*).

**Figure A1:**
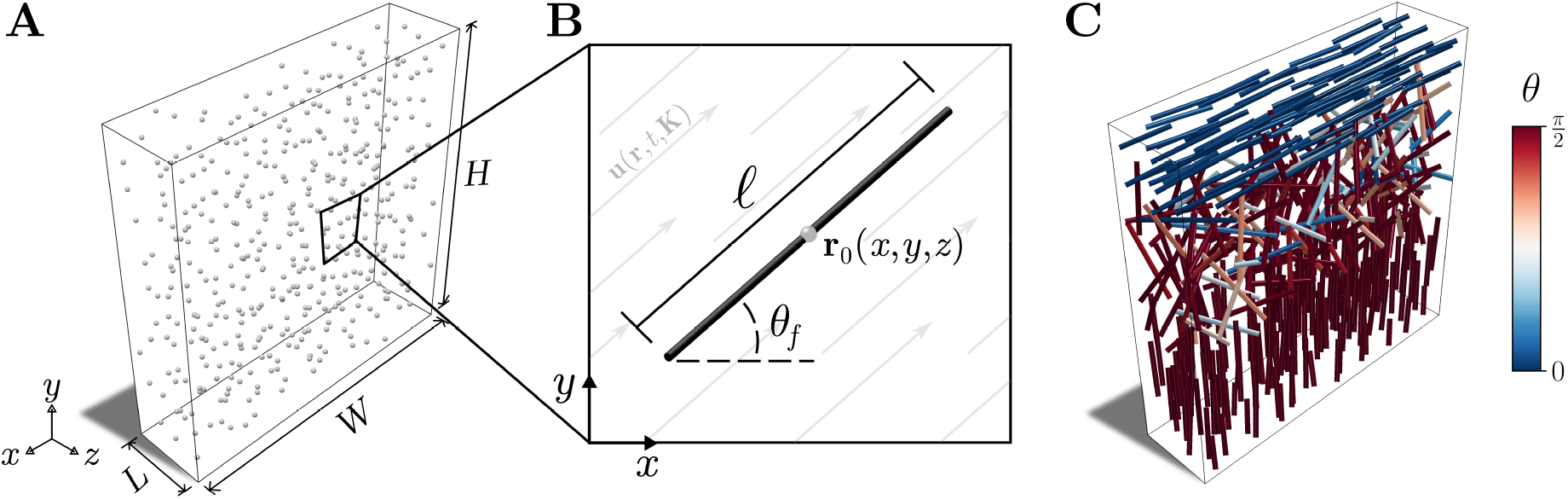
Schematic of discrete fiber model. (A) illustrates the generation of uniformly distributed seed points within the AC domain. (B) shows a 2D schematic of a straight fiber generated through one of the seed points at an orientation *θ*_*f*_. (C) A representative illustration of a three-dimensional discrete fiber network generated through this process.

Further, at each time step, we also remove some of the pre-existing fibers from the system in the following manner. For each fiber we generate a random number *ν* ~ 𝒰(0, 1) and remove the fibers with *ν >* 1 ™ *β*. Here *β* represents the rate of fiber removal and can be dependent on the physiological state of AC. After the deposition and removal of fibers at each simulation step, we calculate the depth-dependent fiber orientation distribution. This distribution is subsequently used to update the permeability tensor **K** at different AC depths using equation (5), which is then use to calculate the local fluid velocity **u**(**r**, *t*, **K**) using equation (3). At each simulation step, we also check if the distribution has reached the steady state (relative to that from the previous steps) to terminate the simulation. The summary of the whole process is shown as a flowchart in Fig. 1C.

## Appendix B Estimation of elastic moduli of AC

In this appendix, we outline the approach used for the extraction of elastic moduli of collagen fiber reinforced AC. A more detailed description of the numerical framework, including implementation details and validation, will be presented elsewhere. Discrete modeling of fibrous networks has been used extensively to study the biomechanics of collagen networks or the extracellular matrix (ECM) (Dong et al., 2017; Filla et al., 2023; Mech and Rizvi, 2024). While collagen fibers serve as the primary load-bearing elements, other components, such as aggregating proteoglycans within the ECM, also play important roles in resisting compression and shear forces. Despite this, most existing studies overlook the mechanical contributions of these non-fibrillar components. To address this, our model of articular cartilage incorporates the non-fibrillar components as a background matrix in which the fibers are embedded. For this, we employ the lattice spring model (LSM), a method commonly used to represent isotropic linear elastic materials (Buerzle and Mazza, 2013; Kakaletsis et al., 2023). The lattice consists of harmonic springs between the nearest regularly spaced nodes and the body-diagonal nodes, forming a cubic network structure. The energy associated with these springs is given by,

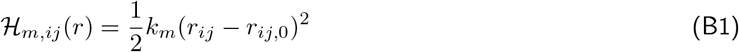

where *k*_*m*_ is the stiffness constant of the lattice.

Next, the fibrillar component of the articular cartilage, that is, collagen fibers, is modeled as semiflexible athermal chains. We have assumed that the fibers have only two modes of deformation - stretching or compression and bending. The energy associated with the stretching or compression of fiber segments is given by

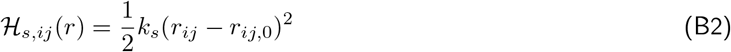

where *k*_*s*_ is the stretching stiffness equal to the product of Young’s modulus *E* and the cross-sectional area *A* of the collagen fiber. *r*_*ij*,0_ and *r*_*ij*_ are the resting and deformed lengths of the fiber segment. The energy associated with the deflection or bending between the fiber segments is given by,

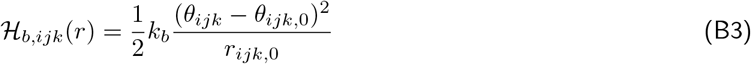

where *k*_*b*_ is the bending stiffness equal to *EI* and *I* is the second moment of area of the fiber. *θ*_*ijk*,0_ and *θ*_*ijk*_ are the initial and deflection angles between the neighboring fiber segments. The equilibrium configuration of the AC is obtained by minimizing the total energy contribution coming from the aforementioned three sources.

To investigate the mechanical response of the articular cartilage, we employed a displacement-controlled approach for compression and shear. Displacement *δ* is applied to the vertices lying on the top face of the network, while the vertices on the bottom face are kept fixed, as shown in Fig. S1(A and D). After applying the displacement, the energy of the system is minimized using the fourth-order Adams-Bashforth method, which offers a good balance between computational efficiency and accuracy, especially for stiff systems of equations. Once the system reaches equilibrium, the force is calculated at the top end of the AC sample, which is processed to extract compressive and shear moduli.

## Appendix C Estimation of effective viscosity of the synovial fluid

To estimate the effective viscosity of the synovial fluid in the presence of articular cartilage, we consider one half of the articular joint, as illustrated in Fig. 1A. In this configuration, the top surface of the AC is taken to represent the centerline of the synovial cavity. We model this system as a Couette flow, wherein a virtual top plate moves at a constant velocity *u*_0_, while the bottom plate, representing the cartilage–bone interface, remains fixed. In the absence of a porous cartilage layer, the effective viscosity experienced by the moving plate would be identical to that of the synovial fluid. However, in the present setup, where the AC includes a collagen fiber network with varying orientation and density, the effective viscosity is modified by the internal flow resistance of the tissue and is therefore expected to deviate from that of the pure synovial fluid.

We solve steady state versions of equations (1)-(3) along with the previously described interfacial conditions to obtain the steady state velocity profile and subsequently shear stress on the top plate, *τ*_steady_. By equating the shear stress obtained here with that in the absence of the AC, we calculate the effective viscosity as

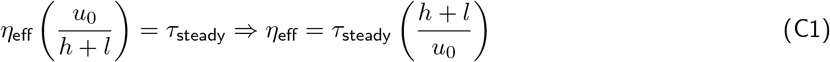

