## Supplementary Figure S1 for "Fluid flow induced biomechanical origin of collagen architecture in articular cartilage"

### Supplementary figures

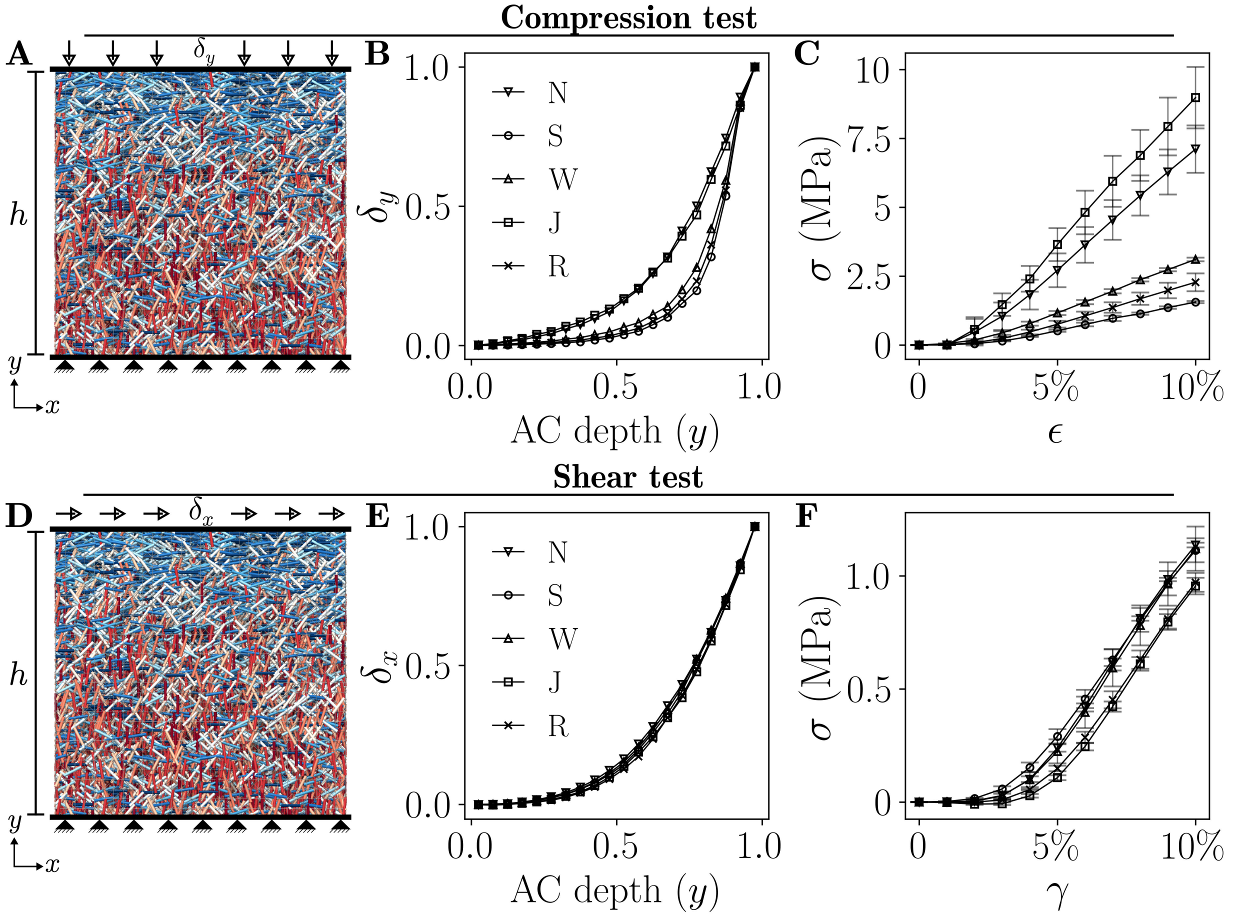


Figure S1: **Mechanical response of articular cartilage under compression and shear**. The figure summarizes the mechanical tests performed on articular cartilages obtained under five distinct physical activities, representing no activity (N), swimming (S), walking (W), jumping (J), and running (R) using a discrete fiber model. (A) and (D) illustrate the schematic of displacement-controlled mechanical loading of cartilage under compression and shear, respectively. The corresponding normalized displacement profile with respect to cartilage depth under (B) compression and (E) shear is shown. The stress-strain plots of the same are shown for (C) compression and (F) shear, averaged over five samples.
